## Supplementary S1-S9 Figures and Supplementary S1-S3 Tables for "*Ustilago maydis* telomere protein Pot1 harbors an extra N-terminal OB fold and regulates homology-directed DNA repair factors in a dichotomous and context-dependent manner"

**S1 — S3 Tables**

### S1 Fig. a

```
metazoan
Chelonoidis_abingdonii            -------MPV--QVID---GTPENLL----PSHLQ-KVD--LKYLQ-MGCDQANKYFQGTVILSYPLTK----------L
Takifugu_rubripes                 -------MPV--HLLSDGAGPPRQVP-----SHLS-RIP--ITLIN-TSVDCTDRTVQGRVVHKGPLVP----------T
Capitella_teleta                  --------MP--FFLT---SEKPDELVNLLPENLR-HLP--LDDIS-NDTETDQIYIRAQISFKGPYQK-----------
Pecten_maximus                    MEKLQCKLCD--IHLK--------IP-----NSLK-KLN--IRDLK-VDTDPSISYIQGRVKERSELQKN----------
Stylophora_pistillata             ----VPIGNG--RYYK---LFAGN-------STPK-RLL--IAELSEPGLL-SSSFIQGIVGQVFPIKSS----------
Orbicella_faveolata               ----FSIGNG--RYCK---LFPGN-------APPR-RVL--IEDLSEPGVL-SSAFIQGTVVKVYPIKCS----------
Strongylocentrotus_purpur         ---MAAFKEM--SESD---LAPEL-------AGLQ-KIS--LSELH-PSSTLLNRYVMGKMVDRSPLIT-----------
Basidiomycetes
Violaceomyces_palustris           ---TNWSKRQ--RILS---EDEEERL-----GRLC-KVDVWLEKVK-QGRG-AAACILGFVLGHRSNLSDRSITLDLSST
Pseudozyma_flocculosa             ---GPAAKRP--RILS---DDDEDRL-----GRLV-DLPRWCERVG-QGRG-AAVCLYATLVKIALSTT-----------
Ustilago_maydis                   ---ATASKRT--RYLN---DADEDRL-----GKLV-SLERWIEGVS-KGRG-AAFCICGHLVQADPSAG-----------
Malassezia_restricta              -----VHPRTPLLALC---ESEVDML-----GAPV-PWSRWLGRVS-RGRG-AAVCTHGTLS--DVSGL----------S
Hydnomerulius_pinastri            ---ENDPKKP--RTDS-----DDDIF-----HESQ-KRSA-KDLPQ-LGID-DSGYIQGKVFMIWPARH-----------
Paxillus_ammoniavirescens         ---GNDAKKP--RTDA-----QDDF------HESR-KRFA-KELPQ-LGKD-DDGYIEGKVFMIWPPRN-----------
Botryobasidium_botryosum          ---DGAAKRP--RIDN---NALDDPF-----DPGQ-ELAP--SELE-VGVSYPSGYVSGVIKIKWPRQG-----------
Suillus_weaverae                  ---DSPPKRP--RLED-----NLSLF-----DETH-RKNA-TDLLD-PATD-ETNNIEAKVHMVFPCVG-----------
Lepista_nuda                      ---GSPSKRP--RVFD----TGELLF-----DDAKQERAA-IDIYR-NGND-DTGYLSGTIFMIWPVMG-----------
Pluteus_cervinus                  ---DEPNKRT--RATE----NVQDEF-----SKLP-EIDV-PELLQ-HGLK-PDRYISAKVFMAWSPMN-----------

consensus                         .....s.tps..+hhs......ts.......sphp.cls..hc.lp..Ghs.ssthlpGplhhhhP.hs...........
```

```
metazoan
Chelonoidis_abingdonii            VNGTEFFKIVLQESENSCS----KTSTINVLFFGRLAG--DCGK----VVHQG--DTMVVA-GFNVA--KSPSANEDNRH
Takifugu_rubripes                 AADSFILKAVVTEESEQ------IVGSINVILQGTLAQ--DFSQ----SVHQG--DVVAAS-GFTVG--KSPTRLRDKLH
Capitella_teleta                  --SRKYFKASLVEG----------NFCINAFFYDKIAE--FCL-----TAAIG--DTIAIS-KMRIE--KNTR-PSQGDH
Pecten_maximus                    -KWFDYMRVSLQDDQDD------TSCQIMVFVTGSIAK--EMD-----IIQKD--YTLVLS-GFKVE--ENSR-IKPSSH
Stylophora_pistillata             ERFGKLCKVVLREDLEKRTRKS-TCATINAFLLGDLAD---EGE----RIKEG--DRVVLS-EVVVE--CSPTD----EH
Orbicella_faveolata               ERFGKLCKILLKEDVGKSSRNS-QLAVISTFLLGELAD---EGS----RICEG--DCVVLA-EVIVE--CSPTG----KH
Strongylocentrotus_purpur         KGKLPVLRLVLEES---------SSTHINVYTFGDFAT---GID----AYKNG--DTIFIS-NAVIE--RSHTFTKDGHH
Basidiomycetes
Violaceomyces_palustris           SSDDRQVNLQLKVIGPN--AAS-LQVYL------QSAG--VL----DRNDRTGLVPSCYVS-GFGGTLAFDSQDASDV--
Pseudozyma_flocculosa             SADGSACHLDLECPPIA-SDPSAGPSKLCAAFYGPLAE--HVFRTLGAVHRRSPPAHVHLS-GFAAT--FDPS------R
Ustilago_maydis                   SVSPDAFNIELSCIHPT--DPS-NRSHLRITFFDQLAS--RVAEPLLLLHRLGPPAIVYLS-GFAAQ-LFDMSAAP----
Malassezia_restricta              QGEPRSFQCVLDGA---------EDAALPVTFRGPYAA--ALHE-VWRRHTHDAPMPVFLS-GFGGD-VVRVRSSPPR--
Hydnomerulius_pinastri            ----GYHRLNLEVVEDS------EIYRFEVEVPNRDGI----------AFRPHERVCLALK-GARSE--KRKESSAPHYL
Paxillus_ammoniavirescens         ----KSHRLNLETVEHS------GIYRFEVEVPHRDGI----------TFRPHQRICLALR-GARAE--RRKESSAPHYL
Botryobasidium_botryosum          --SSKILFHVLNDEE--------PRKTFVVELKGPLVAKYVDFD--RMEVALEDPVLISLKSGEIVK--GNV--GKDKAW
Suillus_weaverae                  --KKRQVNVEMHEHG--------DTYRFLVLISAKVDE--LLPF----PFAVNDRICLSLK-GAQIR--PRPQSSAPCYL
Lepista_nuda                      ----NKLKIIFEARPQ---GEG-GRHKFDIEFTGTCAN--HFQE---LCLKTQDEVQLALK-GATLE--LSG--SQISTL
Pluteus_cervinus                  ----NKHRALVETANAY--SPA-TPHKFEISFSGTCVQ--YFNV---LELKTTDRILILLK-GATLH--KLPKPMSACTL

consensus                         ....phh+l.Lppt.........t.tphpl.h.uphAt...h......thp.s..splhLp.Ghtsc..hssp..t.t.h
```

```
metazoan
Chelonoidis_abingdonii            CCHLEVSEDAG------------------SAVYI--YAKPSIVTASET-
Takifugu_rubripes                 PFNLQLSGDH-------------------ACVFV--SREVPPHPPHPRS
Capitella_teleta                  ACQLIVNND--------------------ACVWA--CVEKPPLHNAPV-
Pecten_maximus                    PCQLVVDEEMST-----------------PKVWY--IEPEPDLIDSQE-
Stylophora_pistillata             QFQLIAWREKS-----------------QASIWV--INNKGNVRANTA-
Orbicella_faveolata               QFQLIVWRQKS-----------------QASIWV--INNKGKVRANTT-
Strongylocentrotus_purpur         KCQLMVHKGRS-----------------HALVWV--CPQQEQGGSNTA-
Basidiomycetes
Violaceomyces_palustris           -PIVDYDQGRIAIKLASGESTSED-------AWF--ERTEHEAE-----
Pseudozyma_flocculosa             PERIEFRTGKAVLRFR-PTSVGDGPD--TEALWV--ERLDRPGN-----
Ustilago_maydis                   -PRVVFAQANAVVKMS-HQSLSADPTAPTSVIWI--EQLATPSA-----
Malassezia_restricta              -AGLVFSERVALKALL-HTDAE-----EELLLWF--EGDRSRPT-----
Hydnomerulius_pinastri            PVTLKYPEG-VVLKYL-SGAN----QGKVFNTWE--GNTHADEW-----
Paxillus_ammoniavirescens         PIILKYPDG-VVLKYL-SGVD----QGKVINTWE--GTT--DEWYN---
Botryobasidium_botryosum          PFKIVFPTE-FVIKFP-ATHRRAELTVDLFNVKEKMGETWFS-------
Suillus_weaverae                  PVMLVFKDG-IAAMLM-SGPS----AGKVFSTWE--GVDWYSST-----
Lepista_nuda                      KMKLLYTEG-ALVKLV-KRRGDLHPINKIINTWI--REKHPSPS-----
Pluteus_cervinus                  PMKLCYEEG-VVLKFL-KRREGIKYDDQTIDTWK--LKEAESNG-----

consensus                         .hpLhhtps.s..................hssWh..tppt.s.t.....
```

### S1 Fig. b

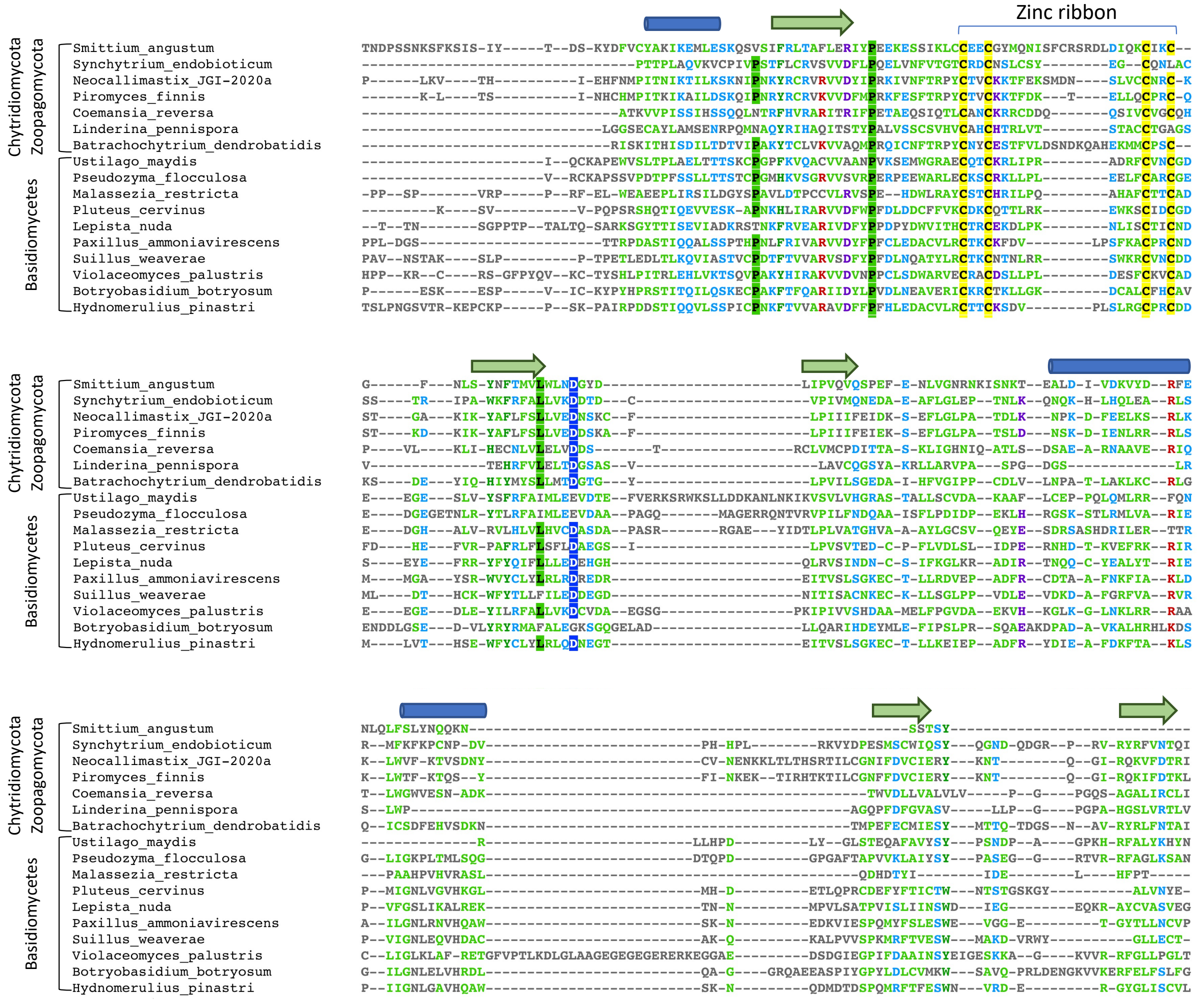
Zinc ribbon
Chytridiomycota
Zoopagomycota
Basidiomycetes
Smittium_angustum
Synchytrium_endobioticum
Neocallimastix_JGI-2020a
Piromyces_finnis
Coemansia_reversa
Linderina_pennispora
Batrachochytrium_dendrobatidis
Ustilago_maydis
Pseudozyma_flocculosa
Malassezia_restricta
Pluteus_cervinus
Lepista_nuda
Paxillus_ammoniavirescens
Suillus_weaverae
Violaceomyces_palustris
Botryobasidium_botryosum
Hydnomerulius_pinastri

### S1 Fig. c

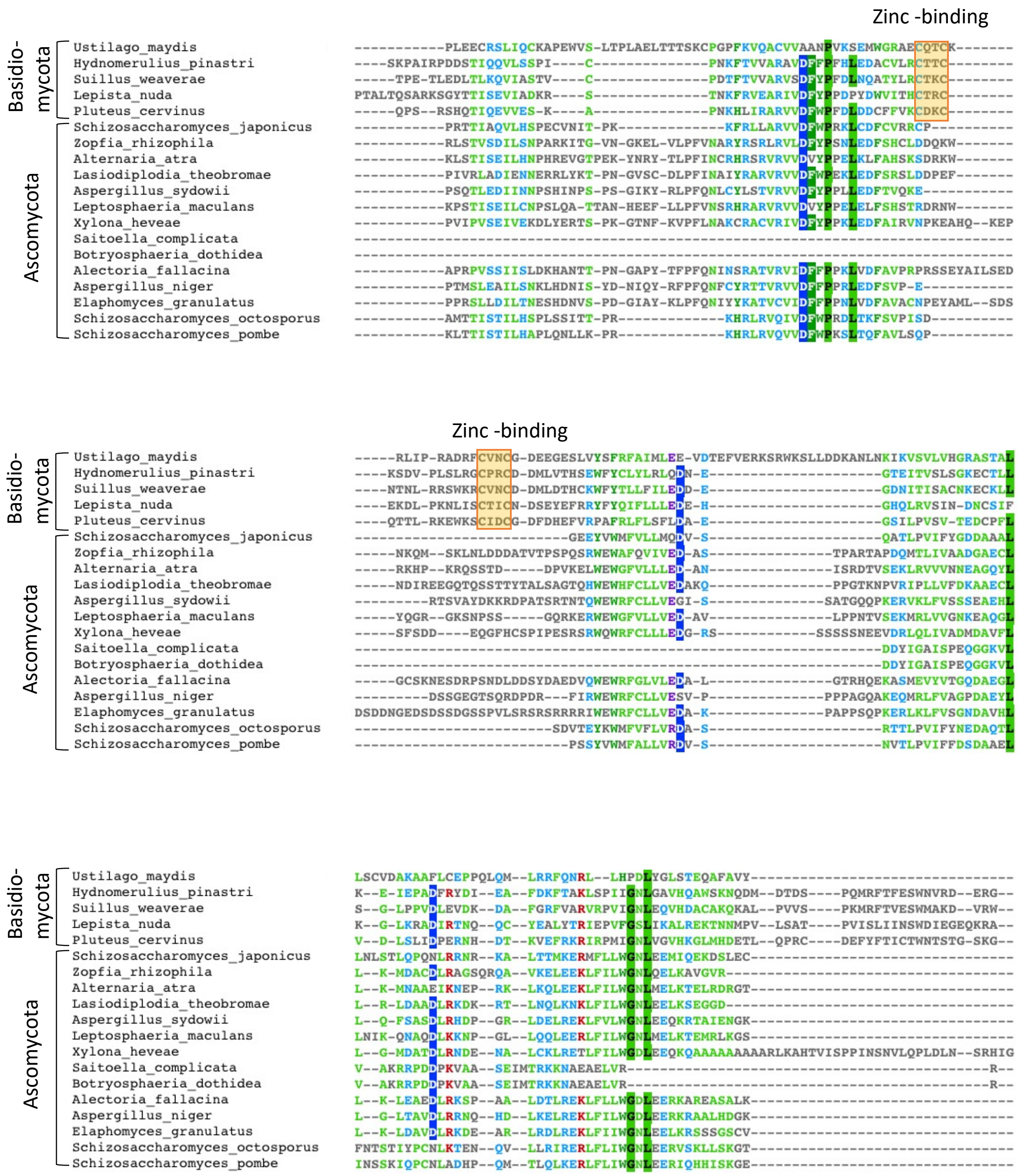

**S1 Fig. Comparisons Pot1 homologs in metazoans and fungi**

**a.** Multiple sequence alignment of the putative OB-N domains from metazoans and basidiomycetes. The alignment and secondary structure predictions were performed using PROMAL3D and displayed using MView. Amino acids were colored by groups.

**b.** Multiple sequence alignment of the OB3 domains of Pot1 homologs from Basidiomycota, Chytridiomycota, and Zoopagomycota.

**c.** Multiple sequence alignment between the OB3 domains of Pot1 homologs from Basidiomycota and Ascomycota. The putative Zinc-binding residues in the basidiomycete OB3 domains are highlighted.

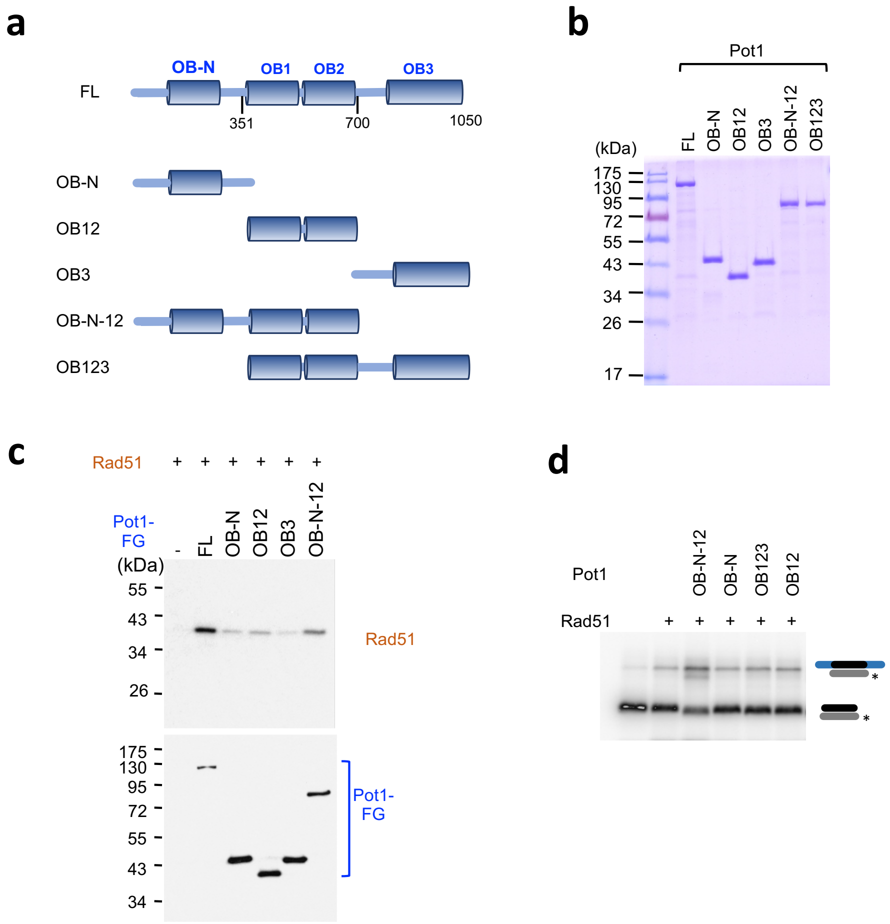

**S2 Fig. The Rad51-binding and stimulatory activities of Pot1 truncation variants**

**a.** The domain structure of Pot1 and the truncation variants analyzed in this study are shown.

**b.** Affinity purified Pot1 proteins were analyzed by SDS-PAGE and Coomassie staining.

**c.** Purified Rad51 was subjected to pull down analysis using FLAG-tagged Pot1 and truncation derivatives. The eluates were analyzed via Western using anti-Rad51 (for Rad51) and anti-FLAG (for Pot1) antibodies.

**d.** The effects of Pot1 truncations on the strand exchange activity of Rad51 were analyzed using oligonucleotide substrates.

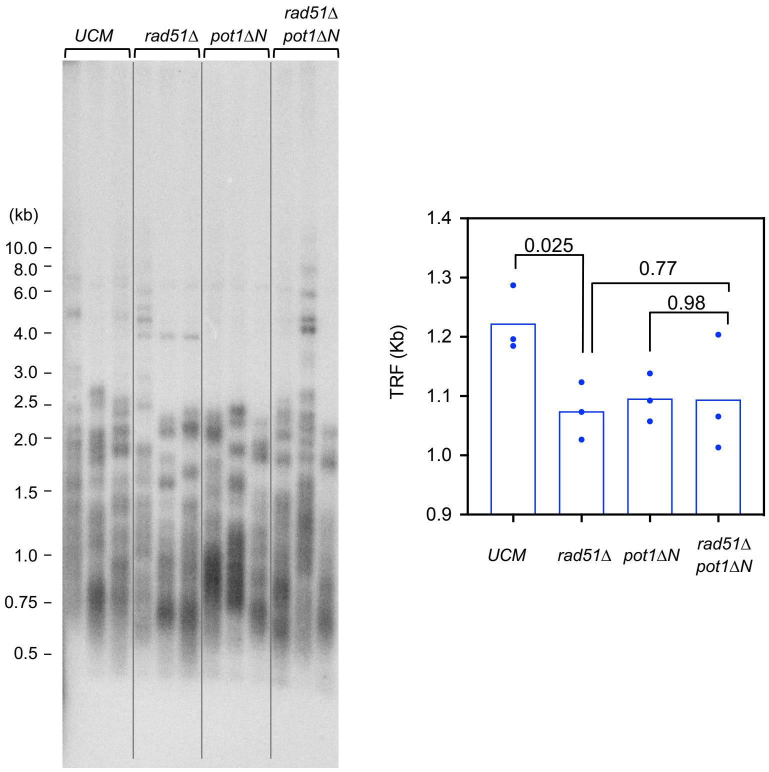

**S3 Fig. Genetic interaction between *rad51* and the OB-N of *pot1***

(Left) Chromosomal DNAs from three independently propagated cultures of each strain were isolated after ~100 generations of growth (4 streaks) and subjected to telomere restriction fragment analysis. (Right) Average TRF lengths were determined and plotted.

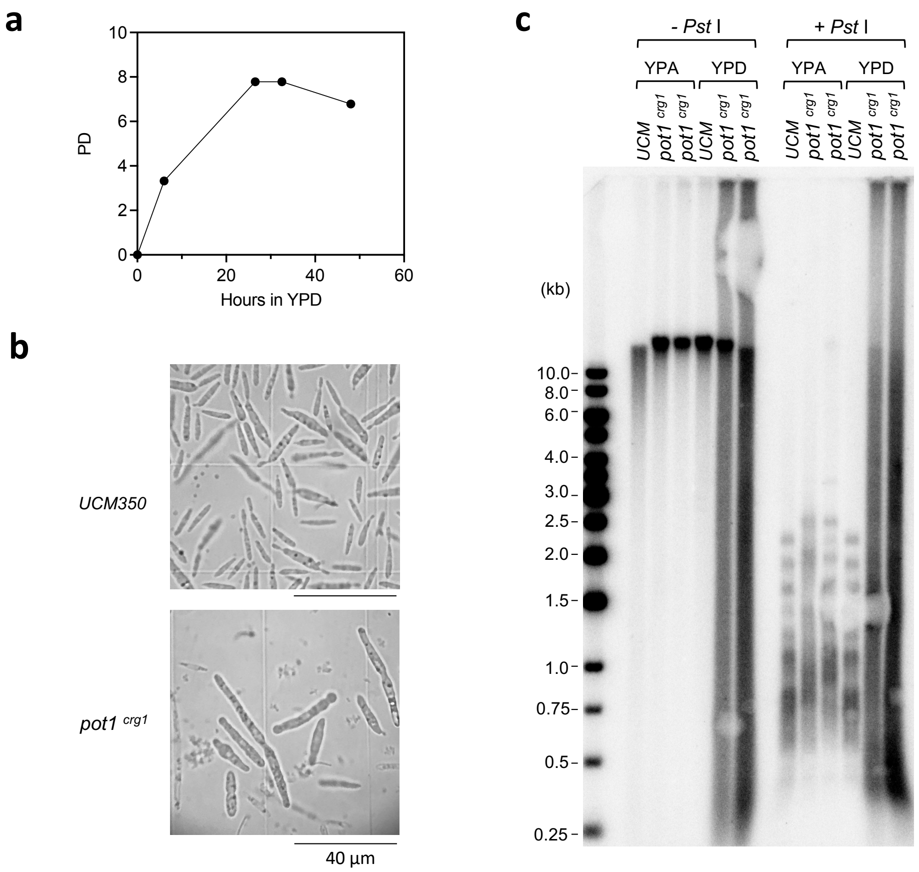

**S4 Fig. Pot1 deficiency impairs cell cycle progression and triggers the production of ECTR**

**a.** A *pot1*$^{crg1}$ liquid culture grown in YPA was harvested, washed 3 times with water, and resuspend in YPD to a starting $OD_{600}$ of 0.1. The population doubling of the culture was then monitored over a period of 48 hours.

**b.** The UCM350 and *pot1*$^{crg1}$ strains were first grown in YPA, and then switched to YPD medium. After another 24 hours of growth, the cells were examined under the microscope.

**c.** Chromosomal DNAs from the indicated strains grown in either YPA or YPD were isolated and subjected to Southern analysis with a telomere repeat probe (TR82) with or without prior *Pst*I digestion.

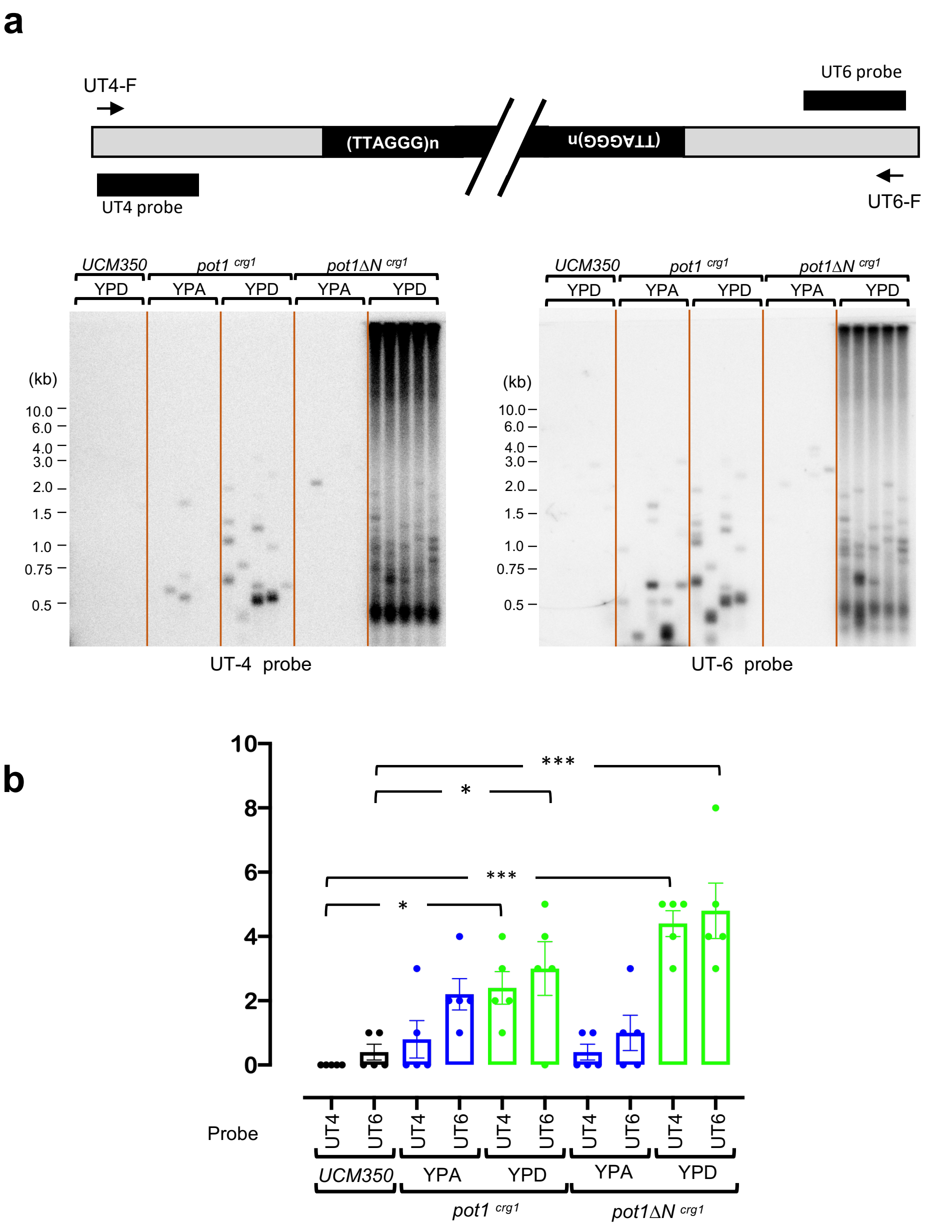

**S5 Fig. The levels of telomere-telomere fusions in *pot1*-deficient *U. maydis***

**a.** (Top) Schematic diagram of the primers and probes used to detect fusions between UT4- and UT6-containing telomeres. (Bottom) Chromosomal DNAs from the indicated strains grown in either YPA or YPD were subjected to PCR-based fusion detection. Five independent PCR reactions per DNA sample were performed; the products were separated by electrophoresis and subjected to Southern analysis using sequentially a UT4 and a UT6 probe.

**b.** The number of fusion fragments were determined and plotted. Statistical significance was calculated using Students' test (*, <0.05; **, <0.01; ***, <0.001).

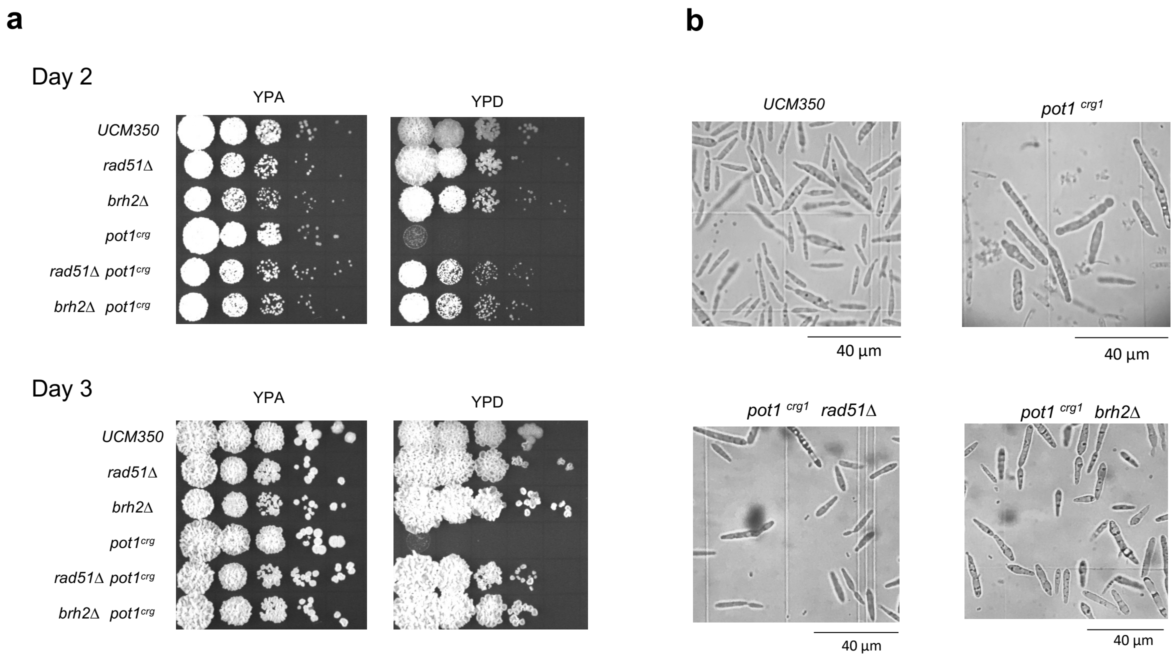

**S6 Fig. The growth of *pot1*, *rad51*, and *brh2* single and double mutants**

**a.** Serial dilutions of the indicated strains were spotted on YPA and YPD plates. The growth of the strains after 2 and 3 days of incubation were imaged.

**b.** The morphologies of the indicated strains after 24 hours of growth in YPD were examined.

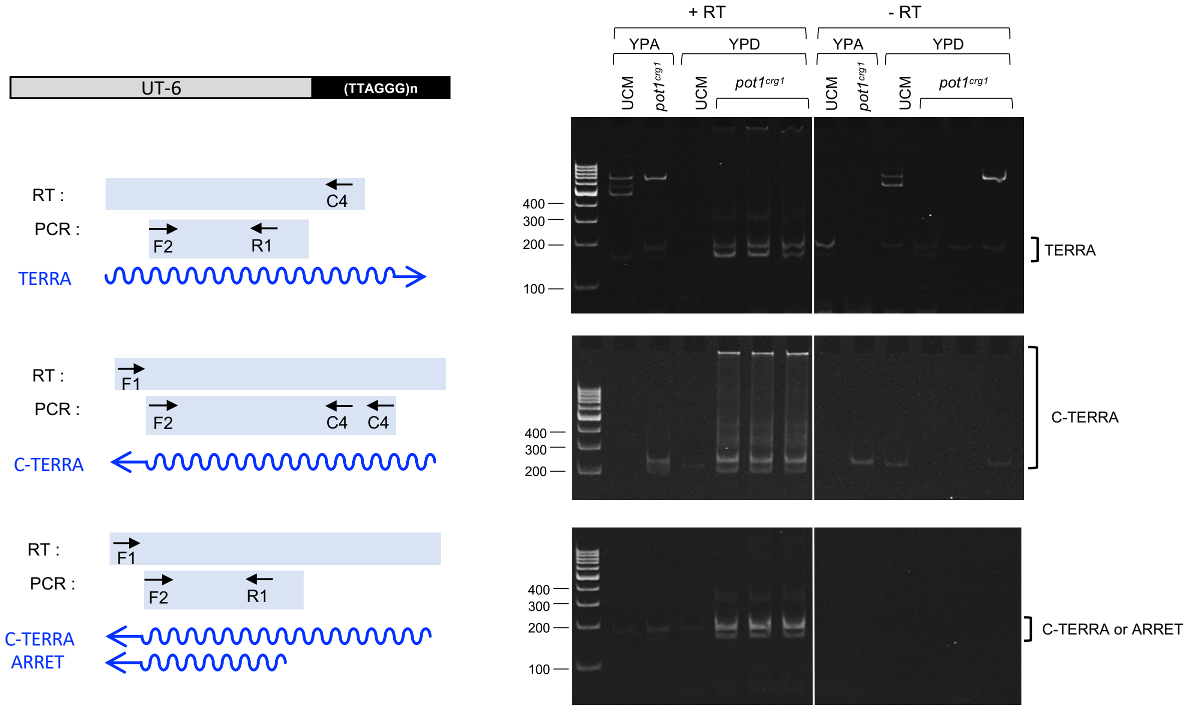

**S7 Fig. Pot1 deficiency triggers the accumulation of telomere repeat containing RNAs**

(Left) Schematic illustrations of the structure of UT6 telomeres and the primers used in the RT and PCR reactions designed to detect telomere repeat RNAs. Note that ARRET was previously defined as C-strand RNA comprised of subtelomere sequences only.

(Right) RNA samples from the indicated strains grown in YPA or YPD were subjected to RT-PCR using the primers indicated to the left of each panel. Both the F1/R2 and F2/R1 PCR reactions are expected to generate ~200 bp products. The multiple bands detected are likely due to minor insertions/deletions in different copies of UT6 elements (Swapna et al., 2018).

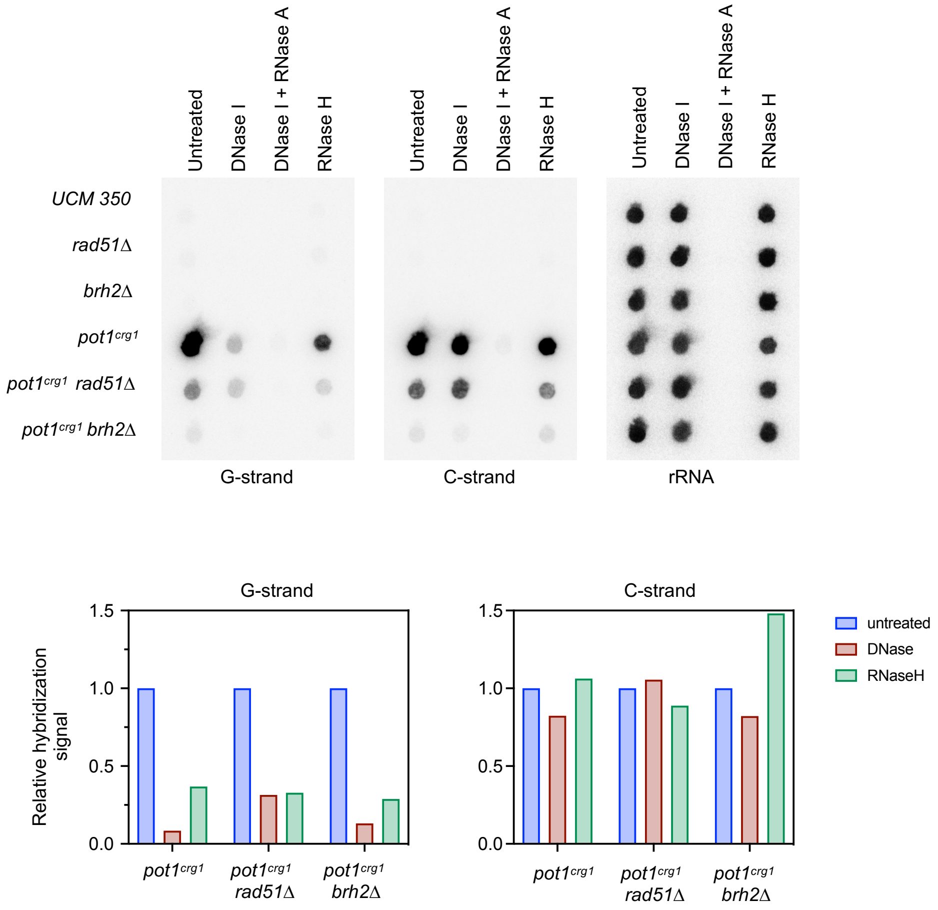

**S8 Fig. The nuclease sensitivity of RNA species from telomeres in pot1-deficient *U. maydis***

(Top) RNAs from the indicated strains grown in YPD were treated with the indicated nucleases and then subjected to dot blot analyses using sequentially probes for detecting G-strand RNA, C-strand RNA, and 26S rRNA.

(Bottom) The ratios of G-strand and C-strand signals to rRNA signals for the untreated, DNase-treated and RNase H-treated samples were calculated. Each ratio was normalized against the untreated sample for the particular strain and plotted.

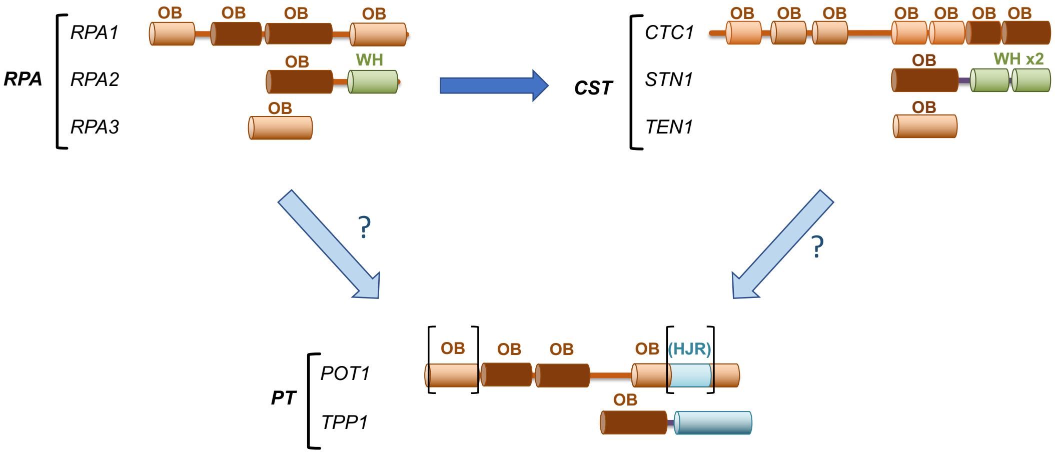

**S9 Fig. Possible evolutionary relationships between three OB-fold rich complexes at telomeres**

The domain structures of RPA, CAT and PT subunits are illustrated schematically. The designations for the domains are as follows: OB, oligosaccharide/oligonucleotide-binding; WH, winged-helix; HRJ, Holliday junction resolvase. The OB folds implicated in ssDNA-binding are shaded in dark brown. Available structural and functional evidence strongly suggests that RPA and CST share a common ancestry (Lue, 2018, Lim et al., 2020). Whether PT is derived from the same primordial ssDNA-binding complex is less clear (Lue, 2018). The demonstration of a 4-OB architecture for Pot1 orthologs in fungi and metazoan suggests that Pot1 and RPA1 have similar domain organizations and may indeed be evolutionarily related.

**S1 Table.** ***U. maydis*** **strains used in this study**

| Alias (Haploids) | *Relevant Genotype* | Reference |
|---|---|---|
| UCM350 [a] | wild type | (Kojic et al., 2002) |
| USZ104 [ab] | *pot1ΔN* | This work |
| USZ105 [ac] | *pot1*$^{crg}$ | This work |
| USZ106 [ad] | *pot1ΔN*$^{crg}$ | This work |
| USZ107 [ae] | *rad51Δ* | This work |
| USZ108 [ace] | *pot1*$^{crg}$ *rad51Δ* | This work |
| USZ109 [af] | *brh2Δ* | This work |
| USZ109 [acf] | *pot1*$^{crg}$ *brh2Δ* | This work |

[a] The genotype of UCM350 *is nar1-6 pan1-1 a1b1*. *nar*, *pan*, and *ab* indicate inability to reduce nitrate, auxotrophic requirement for pantothenate, and mating type loci, respectively.
[b] *pot1* was replaced with an N-terminally truncated allele; a *cbx* cassette expressing the carboxin resistance gene ($Cbx^R$) was inserted upstream of the promoter.
[c] *pot1* promoter was replaced by the *crg1* promoter
[d] *pot1* promoter was replaced by the *crg1* promoter, and the first 1125 nt of the ORF was deleted.
[e] *rad51* was disrupted by a *nat*$^R$-containing cassette.
[f] *brh2* was disrupted by a *nat*$^R$-containing cassette.

**S2 Table. Oligos used in this study**

| Name | Sequence 5' to 3' |
|---|---|
| **Protein Expression** | |
| UmPot1-F-Bam | AAT GGATCC ATG CCG CGC AAG AGA AAA GGT ACC |
| UmPot1-R-FG-NotI | AAT GCGGCCGC CTA CTT GTC ATC GTC ATC CTT GTA ATC CAA TAG ATC GTG TTC GTC AGA T |
| UmPot1-700-R-FG-NotI | AAT GCGGCCGC CTA CTT GTC ATC GTC ATC CTT GTA ATC TCG ATC GGC CAG CGC TTC TTG |
| UmPot1-350-R-FG-NotI | AAT GCGGCCGC CTA CTT GTC ATC GTC ATC CTT GTA ATC AGC TCT GGA TGG ATT TGG TGA A |
| UmPot1-351-F-Bam | AAT GGATCC TTG TCG AGG CAA GCT TCC GCT |
| UmPot1-701-F-Bam | AAT GGATCC CTC GAG CGA CAA TCC AGC CA |
| **Strain Construction and Genotyping** | |
| UmPot1-F-Bam | AAT GGATCC ATG CCG CGC AAG AGA AAA GGT ACC |
| UmPot1-R-FG-NotI | AAT GCGGCCGC CTA CTT GTC ATC GTC ATC CTT GTA ATC CAA TAG ATC GTG TTC GTC AGA T |
| UmPot1-700R-FG-NotI | AAT GCGGCCGC CTA CTT GTC ATC GTC ATC CTT GTA ATC TCG ATC GGC CAG CGC TTC TTG |
| UmPot1-350R-FG-NotI | AAT GCGGCCGC CTA CTT GTC ATC GTC ATC CTT GTA ATC AGC TCT GGA TGG ATT TGG TGA A |
| UmPot1-351F-Bam | AAT GGATCC TTG TCG AGG CAA GCT TCC GCT |
| UmPot1-701F-Bam | AAT GGATCC CTC GAG CGA CAA TCC AGC CA |
| UmPot1(1)Nde | GAAATC CAT ATGCCGCGCAAGAGAAAA |
| UmPot1(700)Xba | ATT TCTAGA AGGAGGAGGCGAAAGCTG |
| UmPot1(-700)Xba | ATA TCTAGA TCGCAAATCGTGACGTGT |
| UmPot1(-1)Eco | ATT GAATTC GATGGATGTGATTCTGGA |
| UmPot1(1126)Nde | GAAATC CAT ATGGGCAACTGCCTCTAC |
| UmPot1(1825)Xba | ATT TCTAGA CGGTGTGGTCGGGATAG |
| Pot1-up-200 | TGACGTGTGACTGACATGC |
| Pot1-1100R (1117-1099) | GCTTGACGTTGGGCGCATT |
| Pot1-1900R (3920-3900) | GTTTTGGATGCGGATGATGTC |
| UmCrg_prom-200 | TGCAACATGAAGTTAGGTGTAGGC |
| Pot1-PCR-1576F | GCCAATTGGTTCTCCGAGCTTCAAG |
| Pot1-PCR-1898R | CCTGTCTTGAGGCCTTCTGCAATGG |
| Pot1-PCR-1801R | GACAGTCTGTGGGCAGATACCTGTC |
| **Strand exchange assays** | |
| Strand_ex_70mer | AGTAGACTCAGCGAACTCACTGATCCAGTCTTAGCATCAGTCACGATACCTCGAGATACATACGG ACGTA |
| Strand_ex_39mer_1 | TGATCCAGTCTTAGCATCAGTCACGATACCTCGAGATAC |
| Strand_ex_39mer_2 | GTATCTCGAGGTCTCGTGACTGATGCTAAGACTGGATCA |

| | |
|---|---|
| **PCR, hybridization, and EMSA assays** | |
| TTAGGG$_4$ (G4) | TTAGGG TTAGGG TTAGGG TTAGGG |
| CCCTAA$_4$ (C4) | CCCTAA CCCTAA CCCTAA CCCTAA |
| TTAGGG$_8$ (G8) | TTAGGG TTAGGG TTAGGG TTAGGG TTAGGG TTAGGG TTAGGG TTAGGG |
| CCCTAA$_8$ (C8) | CCCTAA CCCTAA CCCTAA CCCTAA CCCTAA CCCTAA CCCTAA CCCTAA |
| UmrRNA_26S_121F | GCTTCGGACCATGCCTAAG |
| UmrRNA_26S_642R | CTTGGTCCGTGTTTCAAGACG |
| **RT-PCR** | |
| UT6-TERRA-F1 | GGACGGCAGATATATATTGTGAGTGG |
| UT6-TERRA-F2 | GTGGCAACATTGGGTGAGC |
| UT6-TERRA-R1 | CCGTTGACACATTCAATCCCTC |
| UT6-TERRA-R2 | CTTCAAGCCCTGCAGCC |
| CCCTAA$_4$ (C4) | CCCTAA CCCTAA CCCTAA CCCTAA |
| **STELA and fusion assays** | |
| UT4-F | TCGGGCAACGTTCCATGTCG |
| UT4-subtel-R2375 | CCCTCGAAGGCAGTGCATAC |
| UT6-F | CTACTACACATCGGTTCAGGC |
| UT6-subtel-R2400 | ATGCCAAAGTGGAAATCGTGCAC |
| C Telorette 1 | GCTCCGTGCATCTGGCATCCCCTAAC |
| C Telorette 2 | GCTCCGTGCATCTGGCATCTAACCCT |
| C Telorette 3 | GCTCCGTGCATCTGGCATCCCTAACC |
| C Telorette 4 | GCTCCGTGCATCTGGCATCCTAACCC |
| C Telorette 5 | GCTCCGTGCATCTGGCATCAACCCTA |
| C Telorette 6 | GCTCCGTGCATCTGGCATCACCCTAA |
| Teltail | GCTCCGTGCATCTGGCATC |

**S3 Table. Pot1 homologs utilized for *in silico* analysis in this study**

| Classification | Organism | Protein ID |
|---|---|---|
| Basidiomycota | *Ustilago maydis* | XP_011388186 |
| | *Pseudozyma flocculosa* | SPO37959 |
| | *Malassezia restricta* | XP_027482707 |
| | *Lepista nuda* | KAF9469263 |
| | *Suillus weaverae* | KAG2350098 |
| | *Hydnomerulius pinastri* | KIJ69117 |
| | *Paxillus ammoniavirescens* | KAF8844831 |
| | *Pluteus cervinus* | TFK76258 |
| | *Violaceomyces palustris* | PWN51691 |
| | *Botryobasidium botryosum* | KDQ17907 |
| Ascomycota | *Schizosaccharomyces japonicus* | XP_002173406 |
| | *Zopfia rhizophila* | KAF2190160 |
| | *Alternaria atra* | XP_043173166 |
| | *Lasiodiplodia theobromae* | KAB2573190 |
| | *Aspergillus sydowii* | XP_040708839 |
| | *Leptosphaeria maculans* | XP_003840098 |
| | *Xylona heveae* | XP_018192102 |
| | *Saitoella complicata* | GAO49222 |
| | *Botryosphaeria dothidea* | KAF4308573 |
| | *Alectoria fallacina* | CAF9934687 |
| | *Aspergillus niger* | GAQ40352 |
| | *Elaphomyces granulatus* | OXV06789 |
| | *Schizosaccharomyces octosporus* | XP_013016945 |
| | *Schizosaccharomyces pombe* | NP_594453 |
| Chytridiomycota | *Batrachochytrium dendrobatidis* | XP_006682830 |
| | *Synchytrium endobioticum* | TPX41259 |
| | *Neocallimastix JGI-2020a* | KAG4088494 |
| | *Piromyces finnis* | ORX40370 |
| Zoopagomycota | *Coemansia reversa* | PIA18514 |
| | *Linderina pennispora* | XP_040739463 |
| | *Smittium angustum* | PVZ97799 |
| Metazoan | *Chelonoidis abingdonii* | XP_032656845 |
| | *Takifugu rubripes* | XP_011605395 |
| | *Capitella teleta* | ELU00652 |
| | *Pecten maximus* | XP_033744930 |
| | *Stylophora pistillata* | XP_022798139 |
| | *Orbicella faveolata* | XP_020608671 |
| | *Strongylocentrotus purpuratus* | XP_030828861 |

**Supporting Information References**

KOJIC, M., KOSTRUB, C. F., BUCHMAN, A. R. & HOLLOMAN, W. K. 2002. BRCA2 homolog required for proficiency in DNA repair, recombination, and genome stability in Ustilago maydis. *Mol Cell,* 10**,** 683-91.

LIM, C. J., BARBOUR, A. T., ZAUG, A. J., GOODRICH, K. J., MCKAY, A. E., WUTTKE, D. S. & CECH, T. R. 2020. The structure of human CST reveals a decameric assembly bound to telomeric DNA. *Science,* 368**,** 1081-1085.

LUE, N. F. 2018. Evolving Linear Chromosomes and Telomeres: A C-Strand-Centric View. *Trends Biochem Sci,* 43**,** 314-326.

SWAPNA, G., YU, E. Y. & LUE, N. F. 2018. Single telomere length analysis in Ustilago maydis, a high-resolution tool for examining fungal telomere length distribution and C-strand 5'-end processing. *Microb Cell,* 5**,** 393-403.
